# Structure and function of a hexameric cyanophycin synthetase 2

**DOI:** 10.1101/2023.04.15.537035

**Authors:** Linda M. D. Markus, Itai Sharon, Kim Munro, Marcel Grogg, Donald Hilvert, Mike Strauss, T. Martin Schmeing

## Abstract

Cyanophycin is a natural polymer composed of a poly-aspartate backbone with arginine attached to each of the aspartate sidechains. Produced by a wide range of bacteria, which mainly use it as a store of fixed nitrogen, it has many promising industrial applications. Cyanophycin can be synthesized from the amino acids Asp and Arg by the widespread cyanophycin synthetase 1 (CphA1), or from the dipeptide β-Asp-Arg by the cyanobacterial enzyme cyanophycin synthetase 2 (CphA2). CphA2 enzymes display a range of oligomeric states, from dimers to dodecamers. Recently, the crystal structure of a CphA2 dimer was solved but could not be obtained in complex with substrate. Here, we report cryo-EM structures of the hexameric CphA2 from *Stanieria* sp. at ~2.8 Å resolution, both with and without ATP and cyanophycin. The structures show a trimer-of-dimers hexameric architecture, and substrate-binding interactions that are similar to those of CphA1. Mutagenesis experiments demonstrate the importance of several conserved substrate-binding residues. We also find that a Q416A/R528G double mutation prevents hexamer formation and use this double mutant to show that hexamerization augments the rate of cyanophycin synthesis. Together, these results increase our mechanistic understanding of how an interesting green polymer is biosynthesized.

## Introduction

Cyanophycin is a natural polymer produced by a wide range of bacteria (Sharon et al. 2021; Sharon et al. 2023a). Discovered in cyanobacteria over a century ago (Borzi 1886-1887), it is composed of a poly-L-aspartate backbone with arginine residues covalently attached to the aspartate sidechains (Allen and Weathers 1980) (Figure 1A). Cyanophycin has a nitrogen content of 24% by mass, higher than any other common natural organic polymer. It is insoluble at physiological pH, spontaneously precipitating into membrane-less granules (Dembinska and Allen 1988). These properties, along with its inert nature, make cyanophycin an ideal molecule for storing fixed nitrogen in cells. Various cyanobacteria display different cyanophycin metabolism schemes that are advantageous for the species. For example, some unicellular, diazotrophic species exhibit a day / night metabolic cycle, where excess fixed nitrogen is funneled into cyanophycin biosynthesis during dark hours. These species break down cyanophycin and mobilize the nitrogen for other biosynthetic processes in the daytime when nitrogen cannot be fixed in the cell because the production of molecular oxygen by photosynthesis would inhibit nitrogenase (Li et al. 2001). In cyanobacteria that differentiate into vegetative cells and heterocysts, nitrogen fixation and photosynthesis are separated spatially, as is cyanophycin metabolism: Anaerobic heterocysts accumulate cyanophycin, which, when required, is broken down to dipeptides that are shuttled to photosynthetic vegetative cells to use the constituent nitrogen (Sherman et al. 2000; Popa et al. 2007; Whitton and Potts 2012; Burnat et al. 2014). Cyanophycin can also be used for longer term storage, allowing cyanobacteria to accumulate nitrogen during periods of high availability for later use in nitrogen-limiting periods, including in nitrogen-deficient toxic blooms (Hampel et al. 2019; Lu et al. 2022).

**Figure 1.**
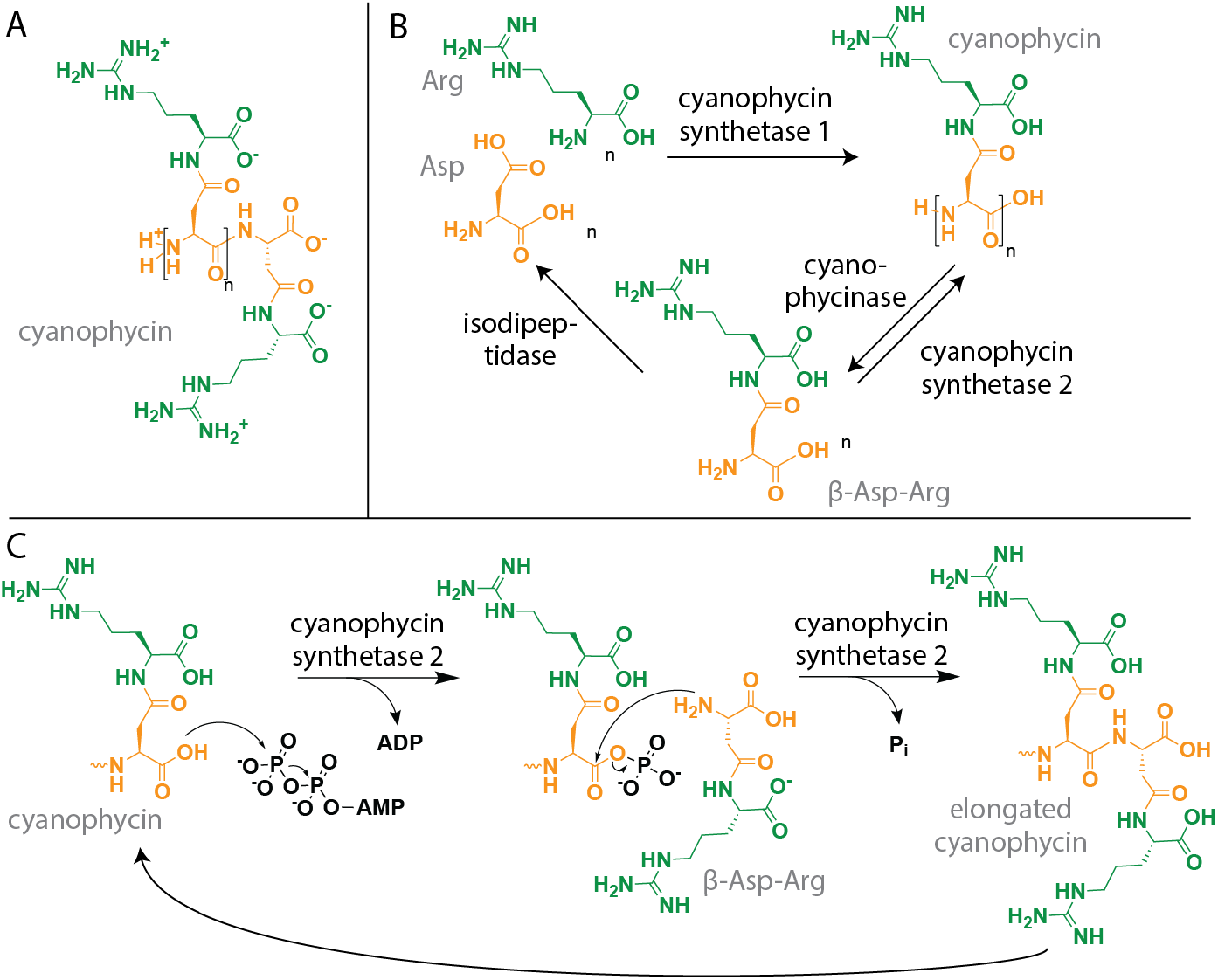
Schematics of cyanophycin and its biosynthesis. (A) The chemical structure of cyanophycin. (B) The known steps of cyanophycin biosynthesis and degradation. (C) The two-step biosynthesis of cyanophycin by CphA2.

In addition to its biological importance, cyanophycin and its derivatives have many potential industrial applications. Cyanophycin itself has been proposed for use in drug delivery (Grogg et al. 2018; Tseng et al. 2018; Grogg et al. 2019), as an active wound dressing (Uddin et al. 2020) and in wastewater treatment (Zou et al. 2022). Alternatively, cyanophycin can be processed to dipeptides for nutritional supplements (Richter et al. 1999; Sallam and Steinbuchel 2010) or to poly-Asp, which is currently used as a biodegradable antiscalant and water softener (Hasson et al. 2011). Accordingly, many studies have focused on increasing the production of cyanophycin in a variety of native and heterologous hosts and growth conditions (reviewed in (Frommeyer et al. 2016)), with a recent analysis showing that attainable yields have already reached commercial viability (Huckauf et al. 2022).

Cyanophycin is biosynthesized by one of two enzymes (Figure 1B,C): cyanophycin synthetase 1 (CphA1) (Ziegler et al. 1998), a common bacterial enzyme that makes it from Asp and Arg in two separate reactions; and cyanophycin synthetase 2 (CphA2) (Klemke et al. 2016), a cyanobacterial enzyme that polymerizes β-Asp-Arg dipeptides. In cells that encode the enzyme, CphA2 is important for balancing cyanophycin biosynthesis and degradation (Picossi et al. 2004) and is under different transcriptional regulation than CphA1 (Picossi et al. 2004). To use the nitrogen, carbon and energy stored in the polymer, cyanophycin must be degraded into its constituent amino acids. This process takes place in two steps: first, a specialized enzyme related to proteases, cyanophycinase, hydrolyses it into β-Asp-Arg dipeptides (Richter et al. 1999); then, these dipeptides are hydrolyzed by one of several enzymes with isoaspartyl dipeptidase activity into free aspartate and arginine (Figure 1B) (Sharon et al. 2023b; Sharon and Schmeing 2023).

CphA2 is, like CphA1, a three-domain enzyme, composed of N, G and M domains (Sharon et al. 2022a). The N domain of CphA2 has no catalytic activity but is important for stability and solubility. The M domain is a truncated, inactive version of CphA1’s catalytic M domain, which is responsible for appending Arg to the Asp side chains, an activity that CphA2 does not require. The G domain has CphA2’s only intact active site, and is very similar to CphA1’s G domain and related to ATP-grasp enzymes. The CphA2 G domain catalyzes the ATP-dependent polymerization of β-Asp-Arg in two steps (Sharon et al. 2021): First, the carboxy-terminus of a nascent cyanophycin chain is phosphorylated by ATP, forming an acyl-phosphate intermediate; then, this intermediate undergoes nucleophilic attack by the incoming β-Asp-Arg, resulting in amide bond formation and extension of the cyanophycin chain by one dipeptide building block (Klemke et al. 2016) (Figure 1C). Our structural and biochemical study of CphA2 suggests that it binds cyanophycin in a similar way to the G domain active site of CphA1 (Sharon et al. 2022a), but no enzyme-substrate co-complex structure is available.

CphA2 enzymes often exist as dimers, with the protomer:protomer interaction involving the G domain (Sharon et al. 2022a), a feature also seen in other ATP-grasp enzymes. Some CphA2 enzymes form higher-order oligomers, including CphA2 from *Stanieria* sp (Sharon et al. 2022a). Since the only CphA2 structure available is that of a dimeric enzyme, it is unclear how higher-order oligomers are assembled and whether such assembly provides catalytic advantages to the enzyme.

Here, we report cryo-EM structures of the hexameric CphA2 from *Stanieria* sp. NIES-3757 (*St*CphA2), both alone and in the presence of ATP and cyanophycin. The structures show how the enzyme binds its substrates and how a CphA2 hexamer is assembled as an open ring-shaped trimer of dimers. Using biochemical and mutagenesis experiments, we identify residues that are important for the enzyme’s catalytic activity and oligomerization. We interrogate the importance of hexamer formation for *St*CphA2 activity by comparing the wildtype hexamer to a mutation-enforced dimer.

## Results

We recently reported that *St*CphA2 exists in solution as a higher-order oligomeric species (Sharon et al. 2022a). To further evaluate this observation and its concentration dependence, we first performed size exclusion chromatography of *St*CphA2 at a range of protein concentrations. At all concentrations assayed (0.9, 1.8, 3.6 mg/ml), the protein migrated as a higher-order oligomer (Supplemental Figure 1A). Analytical ultracentrifugation showed that *St*CphA2 sediments overwhelmingly (94%) as a species of ~434 kDa, close to the mass of a hexamer (428 kDa). A very small fraction of the population (3%) appeared as a dodecamer (observed mass 878 kDa, expected 876 kDa), while no species with lower molecular weights were observed (Figure 2A).

**Figure 2.**
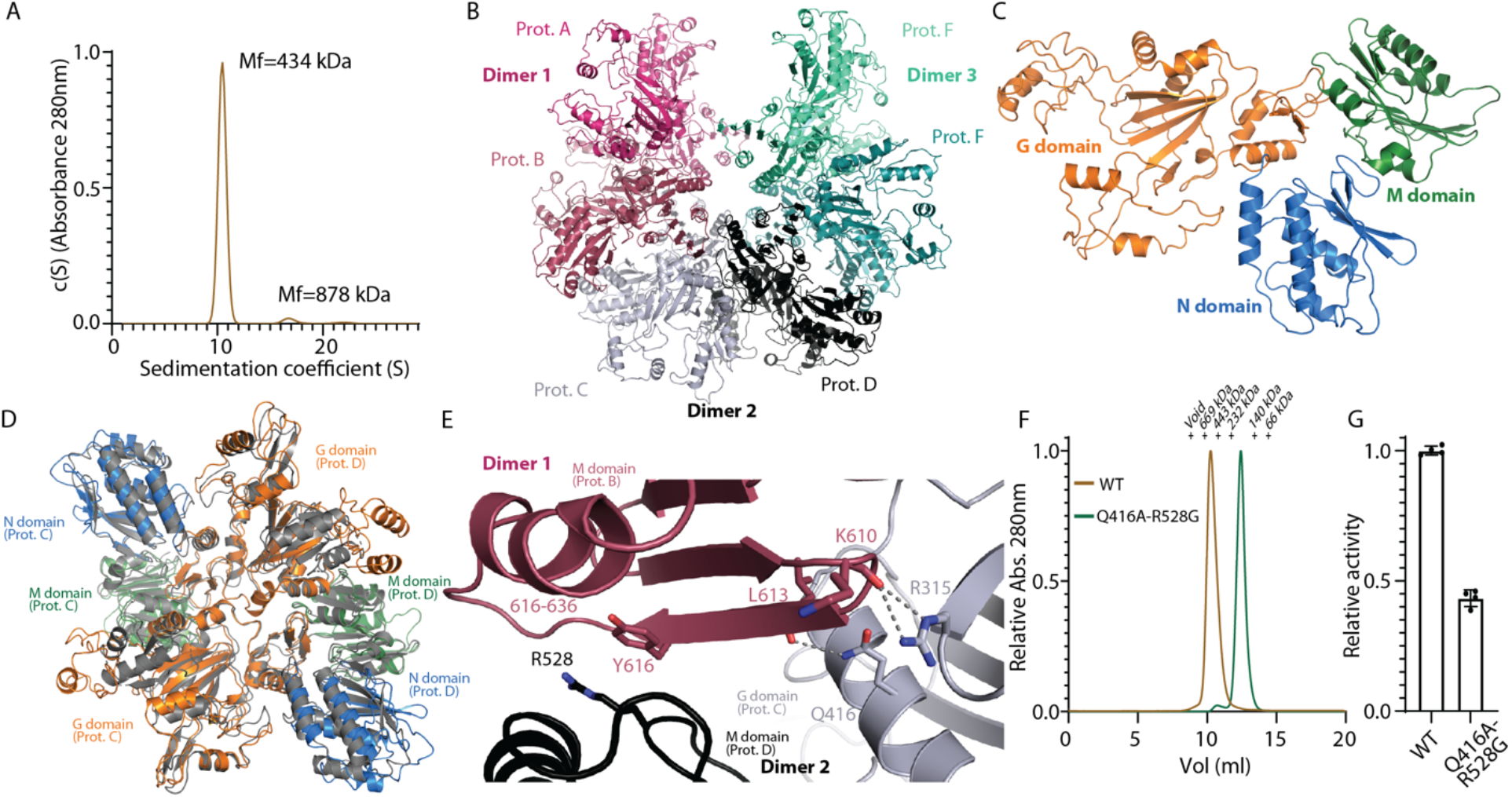
Structure of a hexameric CphA2. (A) Analytical ultracentrifugation analysis of *St*CphA2, showing that the vast majority of the enzyme exists as a hexamer. Quantification shows 94% hexamer, 3% dodecamer and 3% assemblies with a sedimentation value greater than 30 Svedbergs. (B) The cryo-EM structure of hexameric *St*CphA2. The structure is composed of 3 dimers and has C2 symmetry. (C) The structure of a constituent *St*CphA2 protomer, showing the N (blue), G (orange) and M (green) domains. (D) Overlay of the constituent dimer of *St*CphA2 (colored) and *Gc*CphA2 (gray), showing the two enzymes have very similar dimer and monomer architectures. (E) The dimer-dimer interface of *St*CphA2, colored as in panel B. (F) Size exclusion chromatogram analysis of wild type (brown) and *St*CphA2_Q416A-R528G, showing a clear shift in elution volume following disruption of the dimer-dimer interface. (G) Activity rate comparison of wild type (hexamer) and *St*CphA2 Q416A-R528G (dimer).

To understand the nature of *St*CphA2 oligomerization, we determined the enzyme’s hexameric structure using cryo-EM at 2.8 Å (Figure 2, Supplemental Figure 1B,C, Supplemental Table 1). Overall, *St*CphA2 has an architecture best described as trimer-of-dimers that forms an open ring with a small central hole. Each *St*CphA2 protomer has a similar structure to that of the previously published CphA2 from *G. citriformis* (*Gc*CphA2), with an N domain (residues 1-158), G domain (159-486) and M domain (487-636; Figure 2C). In addition, the constituent dimers of *St*CphA2 are also very similar to the dimer of *Gc*CphA2 (Figure 2D). They each have a head-to-head, two-fold symmetric dimer interface involving the G and M domains which, in *St*CphA2, buries 1220 Å^2^ of surface area (compared to 1526 Å^2^ in *Gc*CphA2 (Sharon et al. 2022a)), and the *St*CphA2 and *Gc*CphA2 dimers superimpose with a 2.85 Å RMSD over 854 alpha carbons (which excludes the mobile G_lid_ and G_omega_ subdomains).

The hexameric architecture of *St*CphA2 is assembled from three dimers (Figure 2B). The hexamer has the shape of an open ring with a small central pore. It has an axis of 2-fold symmetry through the gap in the ring, and no 3-fold symmetry that relates the dimers. In the EM maps, the “middle” dimer (protomers C and D) is better ordered than the two peripheral dimers. The interface that oligomerizes the dimers is formed by shape complementarity. The C-terminus of the M domain of protomer B inserts between the G_core_ of protomer C and the M_core_ of protomer D while, reciprocally, the C-terminus of the M domain of protomer C inserts between the G_core_ of protomer B and the M_core_ of protomer A (Figure 2E). Each of these interfaces buries 1684 Å^2^ of surface area. This interaction arranges the dimers into an open ring with angular steps of 108°. The step size does not allow a fourth dimer to be inserted into the ring (which requires steps of ≤90°), and explains the hexameric architecture having 2-fold but not also orthogonal 3-fold symmetry (which requires steps of 120°). The M and G_core_ (sub)domains form the core of the open Ring, with the N, G_lid_ (234-300), and G_omega_ (324-398) (sub)domains protruding outwards (Figure 2B), vaguely reminiscent of the tetrameric CphA1 “spiky ball” architecture (Sharon et al. 2021). The hexameric arrangement of *St*CphA2 orients the G domain active site so it is exposed to bulk solvent, rendering it readily accessible for substrate binding, also reminiscent of CphA1.

To determine the effect of hexamerization on the activity of *St*CphA2, we made a series of structure-guided mutations to the dimer-dimer interfaces. We first deleted the C-terminal 25 amino acids (611-636), which fold into a strand and helix that form interactions with both protomers in an adjacent dimer (Figure 2E). However, the truncation resulted in unstable protein with low solubility. We therefore constructed point mutations that abolish several key dimer-dimer interactions: R315A to eliminate hydrogen bonds with the backbone carbonyl of K610; Q416A to eliminate interactions with the backbone carbonyl of L613; R528G, to eliminate possible interaction with the backbone carbonyl of E617 and van der Waals interactions with Y616; and Y616R, to introduce electrostatic repulsion with R528 (Figure 2E). Gel filtration experiments showed that the point mutants affect oligomerization to varying extents, with *St*CphA2_R528G existing as an approximately 50:50 dimer:hexamer mixture, and *St*CphA2_Q416A as a greater than 75:25 dimer:hexamer mixture (Supplemental Figure 1D). In an attempt to achieve complete dimerization, we produced the Q416A/R528G double mutant.

The combination of these two mutations led to >97% dimer, as assessed by SEC (Figure 2F). We performed circular dichroism (CD) and differential scanning fluorimetry (DSF) with wildtype and Q416A-R528G protein (Supplemental Figure 1E,F). These two enzymes exhibit practically identical CD spectra, suggesting the mutations do not disturb secondary structure. DSF shows Q416A-R528G to have a ~10 °C lower melting temperature (T_m_) than wildtype. The loss of quaternary structure would be expected to lead to a decrease in T_m_. The CD and DSF data indicate that both wildtype and Q416A-R528G should be fully folded under physiological and *in vitro* assay conditions (23 °C). We performed cyanophycin synthesis assays with wildtype *St*CphA2 and the *St*CphA2_Q416A-R528G variant, and found that the dimeric enzyme is around half as active as the native hexamer (Figure 2G).

Since structures of CphA2 in complex with its substrates have not been reported, we collected another cryo-EM dataset of *St*CphA2 in the presence of ATP, (β-Asp-Arg)_4_ and the β-Asp-Arg analog N(2)-succinylarginine (Supplemental Figure 2A, Supplemental Table1). This allowed us to calculate a map with an overall resolution of 2.7 Å, with density for ATP and (β-Asp-Arg)_4_, but not for N(2)-succinylarginine (Figure 3, Supplemental Figure 2B,C, Supplemental Table 1). As expected, ATP binds in the G domain at a position and orientation similar to that seen in *Synechocystis sp*. UTEX2470 (*Su)*CphA1 and other ATP-grasp enzymes. The three C-terminal β-Asp-Arg dipeptides of (β-Asp-Arg)_4_ are visible in the map. As with CphA1, the terminal main-chain carboxylate of cyanophycin is orientated towards the γ-phosphate of ATP. This carboxylate also hydrogen bonds the backbone amide of G455, while reciprocally, the first dipeptide’s backbone amide interacts with the backbone carbonyl of A453. The Arg-guanidino group forms a salt bridge with the side chain of D212 (Figure 3A). The second dipeptide extends towards the N domain and inserts its Arg moiety in a crevice formed by P612 and T163. The third dipeptide has van der Waals interactions with the conserved T202 and hydrogen bonds with the side chain of D212 of the neighboring protomer. The observed interactions between cyanophycin and R308, T163, and D212 (Figure 3A,B) are consistent with the loss of activity seen when the analogous residues in *Gc*CphA2 (*St*CphA2 R308, T163, D212

**Figure 3.**
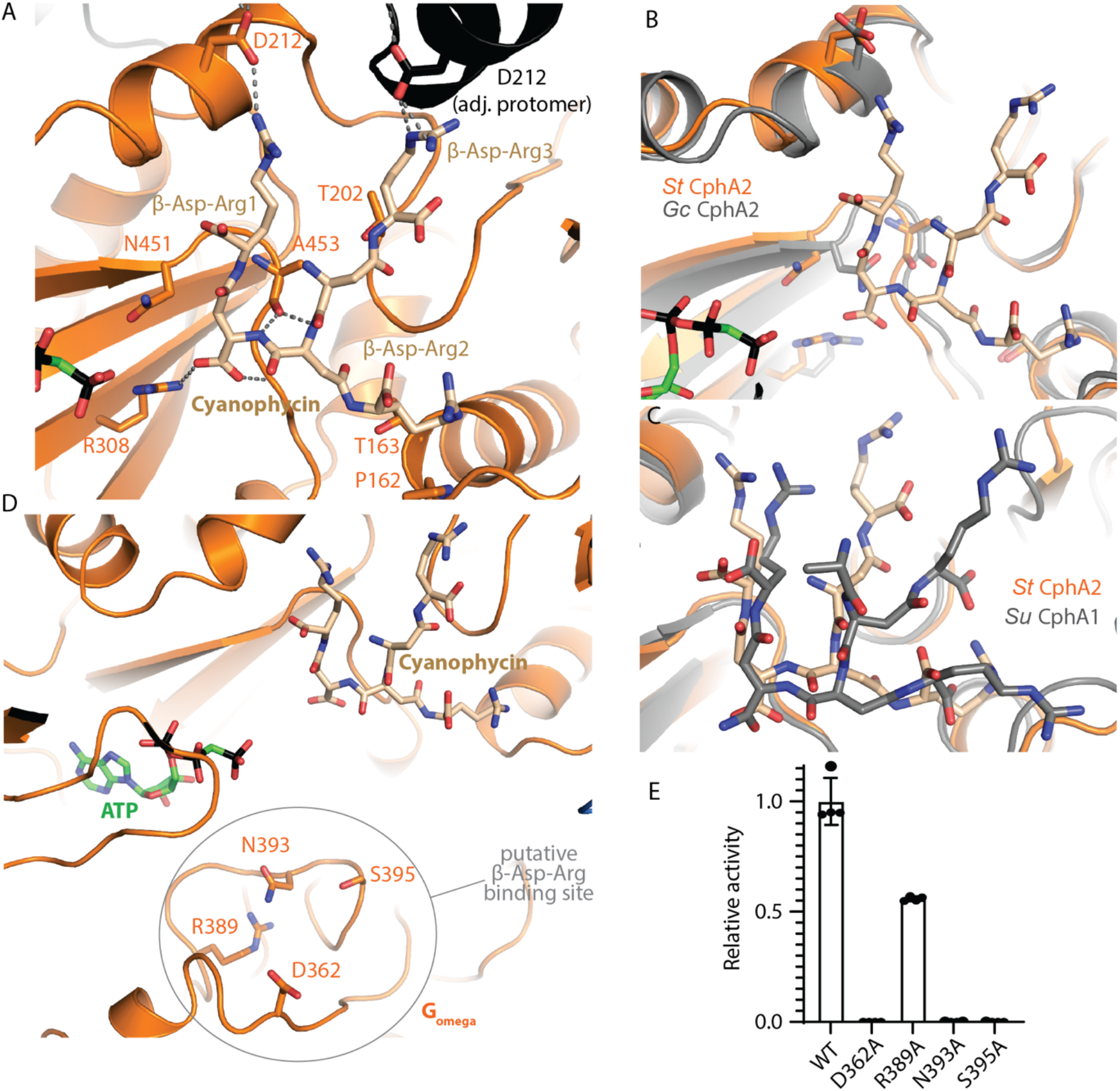
Cyanophycin and ATP binding to CphA2. (A) Binding of cyanophycin to the G domain of *St*CphA2. Key interactions orient the first three dipeptide residues of the nascent cyanophycin chain, allowing its C-terminus to be phosphorylated by ATP. Cyanophycin and ATP were modelled in protomers C and D, but not peripheral dimers AB and EF because the maps are of better quality for the central CD dimer and signal is weaker in the peripheral dimers. (B) Overlay of the G domain active site of *St*CphA2 (orange) and *Gc*CphA2 (gray), showing the two enzymes share high structural similarity around the active site. (C) Overlay of *St*CphA2 (orange) and *Su*CphA1 (gray) G domain active sites, showing a similar binding mechanism and positioning of cyanophycin in the active sites. (D) Several conserved residues around the large loop of G_omega_ that may form the β-Asp-Arg binding site. (E) Mutation of any of these residues to Ala reduce or abolish activity.

≈ *Gc*CphA2 R292, S148, D197) were mutated to alanine (Sharon et al. 2022a). Some details of cyanophycin binding differ between *St*CphA2 and *Su*CphA1. For example, *St*CphA2 H208 interacts with the guanidinium groups of both the first and third dipeptide, but *Su*CphA1 has an isoleucine at the analogous position, likely leading to the small differences in the orientation of the first and third dipeptide we observe (Figure 3C). The G domain loops (*St*CphA2 183-190; *Su*CphA1 181-185) are of different lengths and in different positions in the two enzymes, and there is no direct contact of the loop with bound cyanophycin in *St*CphA2.

Unfortunately, no density could be observed for N(2)-succinylarginine near the large loop of G_omega_, where it is believed to bind (Figure 3D). Data collected in the presence of ADPCP, (β-Asp-Arg)_4_ and the native substrate β-Asp-Arg also showed no density for the latter. Therefore, we attempted to gain insight into β-Asp-Arg binding by assaying alanine mutants of four proximal polar residues (D362, R389, N393, S395) (Figure 3D). This region is conserved among cyanophycin synthetase 2 enzymes (Supplemental Figure 2D), and the chosen residues contain charged or hydrophilic sidechains that are orientated towards the putative binding pocket. The D362A, N393A, and S393A mutants are completely inactive, whereas R389A activity is reduced by ~50% (Figure 3E), supporting the hypothesis that these residues constitute part of the β-Asp-Arg binding site.

Despite the lack of density for the incoming dipeptide, the maps from the dataset that includes substates show the mobile G_lid_ subdomain to be more ordered and in a more “closed” position relative to the apo dataset. Three-dimensional variability analysis shows high flexibility in both datasets, but the average position tilts by ~70° between apo and substrate-containing samples (Supplemental Figure 2E). This is presumably in response to binding of one or multiple substrates. The inability to obtain robust density for the incoming substrate in cyanophycin synthetase G domains is unfortunate but not surprising as it has precedent in other ATP-grasp enzymes, and the mutational data do provide support for the hypothesis that the large loop serves as a binding site for the substrate dipeptide.

## Discussion

This first structure of a higher-order oligomeric CphA2 enzyme displays an elegant open-ring hexameric architecture. Oligomerization is a theme with cyanophycin synthetases (Supplemental Figure 3): CphA1 has been reported as a dimer or a tetramer, but the dimeric form likely arises by disruption of the tetramer because of suboptimal buffer or biophysical conditions (Hai et al. 2002; Arai and Kino 2008; Sharon et al. 2021). Conversely, CphA2 exists typically as a dimer, so the hexamer subject of this study, *St*CphA2, seems to be the exception to the rule (Sharon et al. 2022a).

There is indication that higher-order oligomerization can provide advantages to some cyanophycin synthetase enzymes. Our recent studies of CphA1 examined the influence of the oligomeric state of a CphA1 enzyme which includes a third active site in the N domain responsible for generating primers on biosynthetic rate: Biosynthesis of cyanophycin under conditions where a primer is supplied was the same for tetramers and enforced dimers (Sharon et al. 2021), but that primer-independent activity was faster for tetramers than for enforced dimers (Sharon et al. 2022b). This may be a result of tetramerization exposing substrate cyanophycin chains to higher local concentrations of primer-producing CphA1 N domain active sites. One CphA1 enzyme we studied that does not contain this primer-generating active site also exists as a tetramer, but we could not test whether tetramerization gave it a catalytic advantage because its dimer-dimer interface is so extensive that we could not disrupt its oligomeric state without comprising the enzyme (Sharon et al. 2021).

Similarly, the higher activity of the *St*CphA2 hexamer over the *St*CphA2 dimer may reflect the high local concentration of active sites: because cyanophycin is both a substrate and a product of the reaction, the proximity of nearby active sites could favor rapid re-binding of the elongated cyanophycin polymer. Notably, though, CphA2 enzymes that are naturally dimers exhibit biosynthesis rates in the same range as hexameric *St*CphA2 (Sharon et al. 2022a), rather than that of the enforced-dimeric *St*CphA2_Q416A-R528G. This suggests that subtle modifications to the enzymes, not evident in a superimposition of hexameric *St*CphA2 and dimeric *Gc*CphA2, tune naturally dimeric CphA2 for rapid biosynthesis.

We also report the first substrate co-complex structures of a CphA2 enzyme. That binding of cyanophycin to the CphA2 G domain is very similar to its binding to the CphA1 G domain (Sharon et al. 2021; Miyakawa et al. 2022) is not surprising, since CphA2 evolved from CphA1, and the G domain retains very similar activity. Cyanophycin binding is dominated by electrostatic interactions for both enzymes, which feature similar, shallow active sites (Supplemental Figure 2C), rather than the deep substrate binding pockets seen in some other enzymes. Our inability to capture incoming β-Asp-Arg substrate precludes a comparison of its binding with that of Asp (Sharon et al. 2022b). It is likely a subtle, cryptic change in the G domain large loop that provides specificity for β-Asp-Arg over Asp.

Overall, CphA2 has a simpler task than CphA1, and has become a simpler enzyme. It performs repeated polymerization reactions at a single active site and has dispensed with catalytic activity at the M domain, as well tethering at the N domain which facilitates iterative reactions at G and M domains. This simpler catalytic cycle would not have been an obstacle to the evolution of *St*CphA2 into a hexamer, as higher order oligomerization state can provide high local concentration of the single active sites for cyanophycin recapture, as well as stabilizing quaternary interactions. In any case, the structures and mutagenesis experiments performed here allow contemplation of these different synthetic schemes and further our knowledge of CphA2, an interesting dipeptide polymerase that biosynthesizes a useful green polymer.

## Methods

### Protein expression and purification

For protein expression, plasmids (Sharon et al. 2022a) containing genes encoding *St*CphA2 (WP_096379798.1), *St*CphA2 mutants or *Gc*CphA2 (WP_012598020.1) were used to transform BL21(DE3) *E. coli*. The cells were grown in terrific broth supplemented with 150 μg/ml kanamycin at 37 °C. Upon reaching an OD600 of ~1, the culture temperature was lowered to 18 °C, protein expression was induced by addition of 0.2 mM isopropyl β-d-1-thiogalactopyranoside (IPTG), and the culture was grown for ~20 hours. Cells were harvested by centrifugation at 4000 rpm using a JLA8.1000 rotor for 10 minutes and the cell pellet was resuspended in buffer A (50 mM Tris-HCl pH 7.5, 250 mM NaCl, 3 mM β-mercaptoethanol (BME), 10 mM imidazole) with 100 μM phenylmethylsulfonyl fluoride (PMSF) and a few crystals of DNAseI (Roche) and lysozyme (Bio Basic Inc.). Cells were lysed by sonication and the lysate was clarified by centrifugation at 40,000 g for 30 minutes. The clarified lysate was loaded onto a 5 ml HisTrap HP column (Cytiva) and washed with at least 75 ml buffer A with 30 mM imidazole pH 8. Protein was eluted with ~20 ml buffer B (50 mM Tris-HCl pH 7.5, 250 mM NaCl, 250 mM imidazole pH 8, 3 mM BME) and concentrated to <4 ml using 100,000 kDa MWCO Amicon® Ultra spin concentrators (Millipore). Protein was loaded on a Superdex 200 pg 16/60 column (Cytiva) equilibrated in buffer C (20 mM Tris-HCl pH 7.5, 100 mM NaCl, 1 mM dithiothreitol (DTT)). Fractions with the highest purity were pooled and concentrated.

Aliquots of purified protein were stored in 20% glycerol at −80 °C.

### Site-directed mutagenesis

Site-directed mutagenesis of *St*CphA2 was by performing PCR reactions with 10 ng template in Phusion® HF buffer (NEB), 0.2 mM of each dNTP, 0.5 mM each of forward and reverse primers (Supplemental Table 2) and 0.5 U Phusion® HF DNA polymerase (NEB) in a reaction volume of 25 μl. Following the reaction, template DNA was digested by incubation with DpnI (1 μl; NEB) at 37 °C for 1 hour. Two microlitres of the reaction were used to transform *E. coli* DH-5α cells, which were plated on LB agar plates containing 50 µg/ml kanamycin and incubated overnight at 37 °C. For plasmid preparation, single colonies were cultured in LB medium containing 50 µg/ml kanamycin overnight. Plasmid DNA was isolated using EZ-10 Spin Column Plasmid DNA Miniprep Kit (BioBasic) and sequenced by Sanger sequencing at the Centre d’expertise et de services Génome Québec.

### Size exclusion chromatography and analytical ultracentrifugation

Size exclusion chromatography was performed with an S200 increase 10/300 column (Cytiva) in buffer C at 4 °C, using a flow rate of 0.5 ml/min and injection volume of 150 μl. The column was calibrated with an Amersham™ HMW Calibration Kit.

Sedimentation velocity analysis by AUC was performed using a Beckman Coulter Optima analytical ultracentrifuge. The protein sample was dialyzed extensively against a buffer containing 20 mM Tris-HCl pH 7.5, 100 mM NaCl, 1 mM Tris-(2-carboxyethyl)phosphine (TCEP) and diluted to a concentration of 0.8 mg/ml. The sample and reference buffer were housed in a 12 mm 2-sector Epon-charcoal cell. AUC was performed at 10 °C using an An-60 Ti rotor at 18,000 rpm, and sedimentation behavior was observed by recording 175 absorbance scans (280 nm) at intervals of 180 seconds. The oligomeric distribution was obtained by fitting the resultant data set to the Lamm equation model using the SEDFIT software package and temperature-corrected buffer and protein parameters generated by SEDNTERP.

### Cyanophycin synthesis assays

Cyanophycin synthetase-catalyzed polymerization of β-Asp-Arg (Sharon et al. 2021) was performed in reaction mixtures containing 2 μM purified enzyme, 2 mM ATP, 2 mM β-Asp-Arg, 50 μM synthetic cyanophycin (β-Asp-Arg)_3_ as a primer (Sharon et al. 2022b), 100 mM HEPES pH 8.2, 5 mM MgCl_2_ and 20 mM KCl in a total volume of 100 μl. Reactions were executed at 23 °C in quadruplicate. Formation of cyanophycin was measured by the increase of absorption at 600 nm from light scattering by insoluble cyanophycin, using a SpectraMax Paradigm spectrophotometer (Molecular Devices), with 5 second linear shaking between reads, for ~2 hours. Data were analyzed with GraphPad Prism. For the calculation of activity rates, the maximum of the first derivative of each OD600 curve was used and smoothed with a 2nd order polynomial to reduce noise.

### Cryo-EM

Samples used for structure determination were the following: (1) 2.2 mg/ml *St*CphA2 in 20 mM Tris-HCl pH 8, 100 mM NaCl, 1 mM DTT; (2) 2.4 mg/ml *St*CphA2, 1 mM (β-Asp-Arg)_4_, 20 mM N2-succinyl-Arg, 5 mM ATP in 20 mM Tris-HCl pH 8, 100 mM NaCl, 1 mM DTT, 10 mM MgCl_2_, 10 mM KCl, with a 30-minute incubation period at 20 °C. For vitrification, 3 μl sample was applied to glow-discharged grids, blotted for 1.5 seconds at force 10, and plunge-frozen in liquid ethane using a Vitrobot Mark IV (Thermo Fisher Scientific).

Grids were imaged on a 300 keV Titan Krios G3 Transmission Electron Microscope (Thermo Fisher Scientific) equipped with GIF BioQuantum LS imaging filter (Gatan) and K3 24MP direct electron detector (Gatan). For all datasets, SerialEM was used to collect micrographs of 40 frames at a dose of 2.0 e^−^/Å^2^/frame, pixel size of 0.675 Å/pixel, and defocus range of −1.0 to −2.5 μm. For dataset (1), the dose rate was 17.57 e^−^/pixel/s and 5663 micrographs were collected. For dataset (2), the dose rate was 7.51 e^−^/pixel/s and 8639 micrographs were collected (Supplemental Table 1).

Data processing up to and including Bayesian polishing was performed in RELION4.0 (Scheres 2012). Beam-induced drift was corrected by MotionCor2 and CTF estimation was performed by CTFfind4. Particle picking was performed using Topaz. Datasets (1) and (2) initially included 769,492 and 1,204,400 particles, respectively. Particles were extracted from the micrographs and classified in 2D. Well-ordered classes were selected, with 597,148 and 707,062 particles respectively used for 3D classification. For the dataset without substrate, an *ab initio* model was generated in RELION to use as a reference for 3D classification. For later datasets, the apo model was low-pass filtered to 60 Å and used as a reference. Classes consisting of hexameric particles were selected (251,001 and 549,663) and refined first without, and then with, a mask. Higher-order aberration refinement, anisotropic magnification and per-particle defocus were performed. Two rounds of Bayesian polishing in RELION followed by 3D refinement with CTF refinement in CryoSPARC3 (Punjani et al. 2017) were performed. Local resolution and local filtering were performed in CryoSPARC3. 3D variability was assessed in CryoSPARC3.

Maps obtained by local filtering were subjected to local anisotropic sharpening in Phenix (Adams et al. 2011). For model building, an initial monomeric model was generated from the protein sequence using Alphafold2 (Jumper et al. 2021) and the published structure of *Gc*CphA2 (Sharon et al. 2022a). Six copies of the model were manually fitted into the density in Coot (Emsley and Cowtan 2004). The model was subjected to real-space and rigid-body refinement in Phenix (Adams et al. 2011).

### Differential scanning fluorimetry

DSF assays were performed with 0.7 mg/ml protein in a buffer containing 50 mM Tris-HCl pH 7.5, 100 mM NaCl, 1 mM DTT and 5x SyproTM Orange in a total reaction volume of 20 µl. The temperature was increased from 5 °C to 95 °C over 2 hours and readings taken using a One Step Plus RT-PCR (Applied Biosystems).

### Circular dichroism

CD spectra were collected at 23 °C and a protein concentration of 0.9 mg/ml in a buffer containing 20 mM Tris-HCl pH 7.5, 100 mM NaCl and 1 mM DTT using a Chirascan instrument (Applied biophysics).

## Data availability

The EM maps and atomic coordinates have been deposited into the EMDB with accession codes 29533 and 29534, and into the Protein Data Bank with accession codes 8FXH and 8FXI.

## Acknowledgements

Thank you to the Schmeing, Strauss and Hilvert labs for stimulating discussions, and Kaustuv Basu and Kelly Sears of the McGill Facility of EM Research for facilitating EM data collection. This work was supported by CIHR Project Grant 178084 and a McGill University James McGill Professorship to TMS, funds from the ETH Zurich to DH, and FRQ Programme de bourses d’excellence pour étudiants étrangers (PBEEE) Short-Term Researcher award to LMDM.

## Author contributions

LMDM: investigation (lead), Writing – original draft preparation (lead); IS: investigation (supporting), methodology (lead), writing – review & editing (equal); KM: investigation (supporting); MG: investigation (supporting); DH: writing – review & editing (equal); MS: supervision (equal), writing – review & editing (supporting); TMS: conceptualization (lead), funding acquisition (lead), supervision (equal), writing – review & editing (equal)

## Supplemental Information

Supplemental Tables 1-2, Supplemental Figures 1-3.

