## Supplemental Information for "Structure and function of a hexameric cyanophycin synthetase 2"

### Supplemental Tables

| | *St*CphA2 (EMD-29533; PDB 8FXH) | *St*CphA2 + ATP + (Asp-Arg)$_4$ (EMD-29534; PDB 8FXI) |
|---|---|---|
| **Data collection and processing** | | |
| Magnification (x) | 130,000 | 130,000 |
| Voltage (keV) | 300 | 300 |
| Electron exposure ($e^-$/Å$^2$) | 80 | 80 |
| Defocus range (μm) | -1.0 to -2.5 | -1.0 to -2.5 |
| Pixel size (Å) | 0.675 | 0.675 |
| Symmetry imposed | C2 | C2 |
| Initial particle images (no.) | 769,492 | 1,204,400 |
| Final particle images (no.) | 251,001 | 549,663 |
| Map resolution (Å) | 2.8 | 2.7 |
| FSC threshold 0.143 | | |
| Map resolution range (Å) | 2.4 to 10.3 | 2.3 to 8.4 |
| **Refinement** | | |
| Model resolution (Å) | 2.8 | 2.7 |
| Model composition | | |
| Non-hydrogen atoms | 30228 | 29643 |
| Protein residues | 3786 | 3678 |
| Ligands | 0 | 8 |
| *B* factors (Å$^2$) | | |
| Protein | 155.75 | 137.06 |
| Ligand | | 20.22 |
| R.m.s. deviations | | |
| Bond lengths (Å) | 0.006 | 0.012 |
| Bond angles (°) | 1.571 | 2.193 |
| Validation | | |
| MolProbity score | 1.92 | 2.23 |
| Clash score | 6.73 | 6.73 |
| Poor rotamers (%) | 1.16 | 3.12 |
| Ramachandran plot | | |
| Favored (%) | 91.84 | 92.08 |
| Allowed (%) | 7.79 | 7.07 |
| Disallowed (%) | 0.37 | 0.85 |

**Supplemental Table 1.** Statistics for cryo-EM data processing and structure refinement.

| Primer name | Sequence |
| --- | --- |
| R315A_F | CTGTGTGTTAACGGTGCCTTCGTGGCG |
| R315A_R | CGCCACGAAGGCACCGTTAACACACAG |
| D362A_F | CTGGAGGAACAGGGTCTGGATCTG |
| D362A_R | GTACAGGTGCATAGCTTCAGCAGTACGG |
| R389A_F | CTGTCTAGCGGTGGCTTCAGCATC |
| R389A_R | GTTAGCAACTTTAGCCAGGTAGATGGTACGG |
| N393A_F | GTAAAGTTGCTGCCCTGTCTAGCGGTG |
| N393A_R | GCAGGTAGATGGTACGGTCACGATC |
| S395A_F | CTAACCTGGCTAGCGGTGGCTTCAG |
| S395A_R | CAACTTTACGCAGGTAGATGGTACGGTC |
| Q416A_F | GATAACATTATCCTGGCGGCAGACATCGCGC |
| Q416A_R | GCGCGATGTCTGCCGCCAGGATAATGTTATC |
| R528G_F | GGTATCCTGATTAACGGTTCTGAGAAAATTCTG |
| R528G_R | CAGAATTTTCTCAGAACCGTTAATCAGGATACC |
| G611trunc_F | CCATTAAACGTAAAGGTGAGAATTTGTACTTCC |
| G611trunc_R | GGAAGTACAAATTCTCACCTTTACGTTTAATGG |
| Y616R_F | GCTGGAACAGCGTGAACTGGAAGC |
| Y616R_R | GCTTCCAGTTCACGCTGTTCCAGC |

**Supplemental Table 2.** Primers used in this study.

### Supplemental Figures

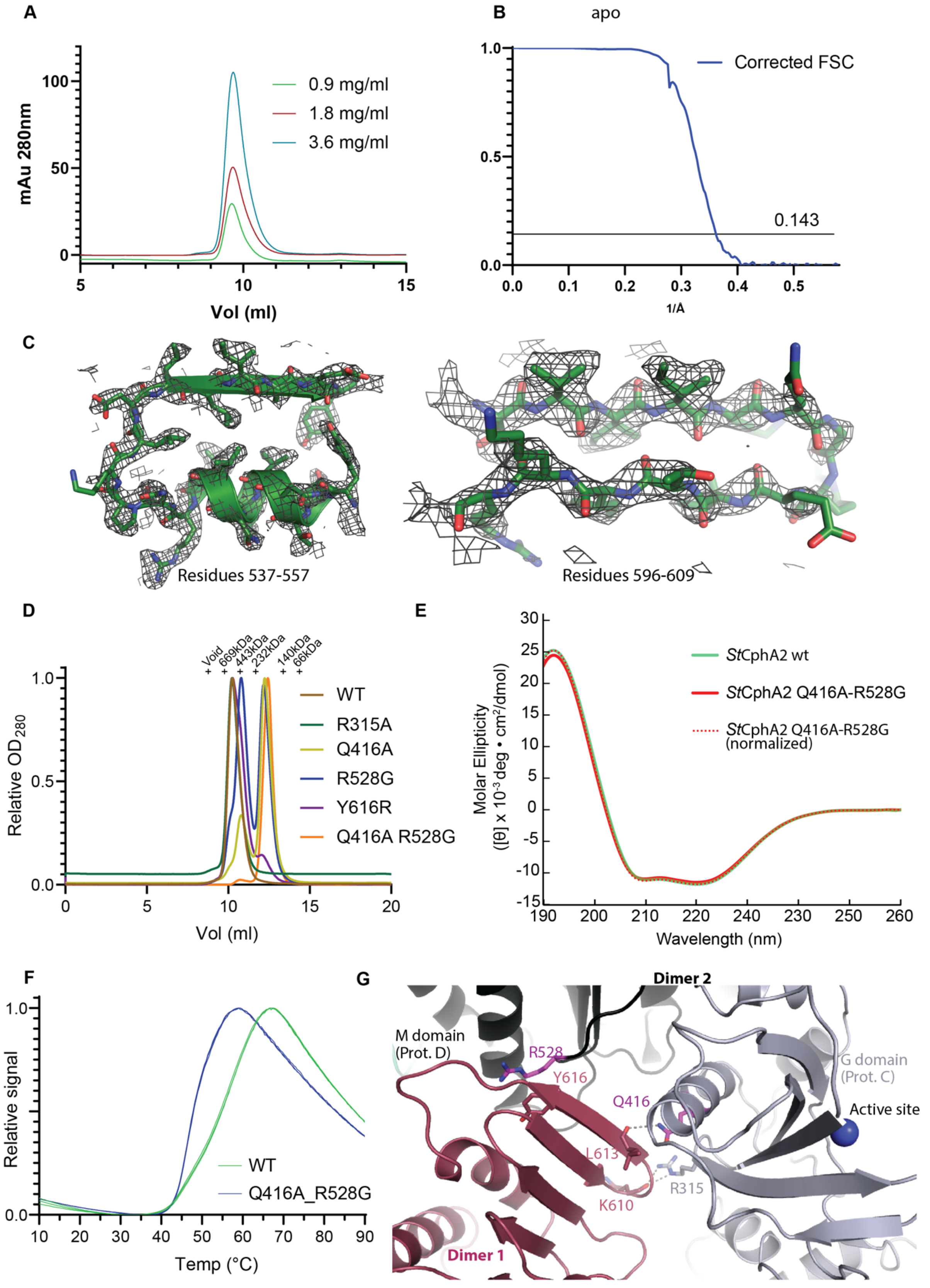

**Supplemental Figure 1.** (A) SEC chromatogram of WT *St*CphA2 at different sample concentrations, showing they all have the same elution volume. (B) FSC curve of apo *St*CphA2 hexamer processed with C2 symmetry. (C) Example of the cryo-EM maps of apo *St*CphA2. The maps are displayed on the left at a contour level of 17 and carve value of 3 Å (residues 537-557) and on the right at a contour level of 22 and a carve value of 3 Å (residues 596-609). (D) SEC chromatogram of *St*CphA2 with different dimer-dimer interface mutations. The mutants display different elution volumes, suggesting the mutations lead to differences in oligomeric state and hexamer stability. (E) Circular dicrhosim spectrum of WT and Q416A R528G *St*CphA2. If the mutant is normalized by multiplication by 1.03 to compensate for 3% error in concentration, the spectra overlay exactly. (F) Differential scanning fluorimetry melting curves of WT and Q416A R528G *St*CphA2. The mutant has a lower $T_m$ than the WT. (G) The active side is ~20 Å from position of mutated Q416, and much further from R528.

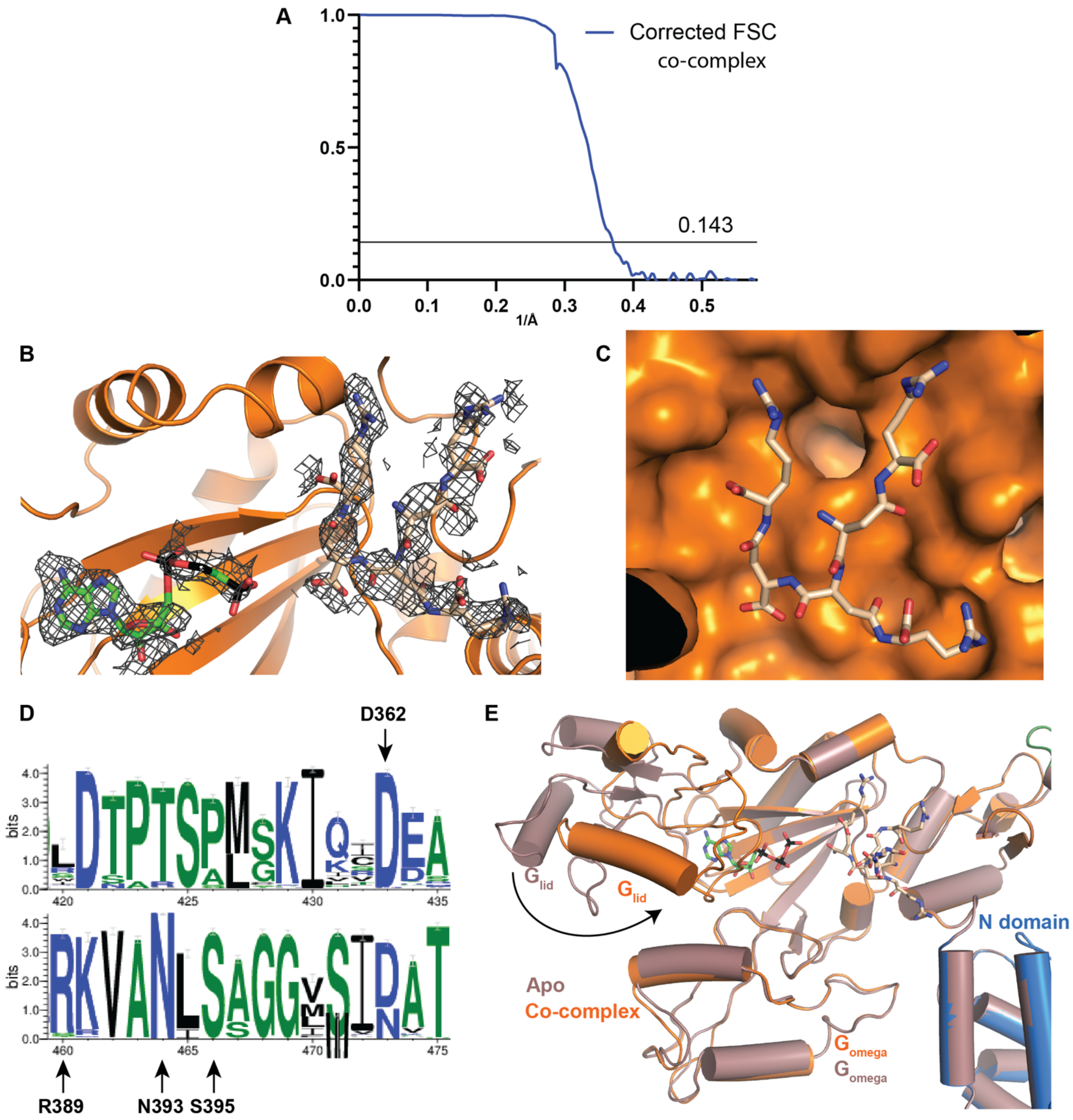

**Supplemental Figure 2.** (A) FSC curve of substrate-bound *St*CphA2 hexamer processed with C2 symmetry. (B) EM density map around AMPPCP and cyanophycin in the active site of *St*CphA2. The map is displayed at a contour level of 5 and carved at 2 Å. (C) Surface rendering of the cyanophycin binding site shows it to be shallow. (D) Weblogo(Crooks et al. 2004) analysis of residues around the large loop close to the G domain active site, showing high sequence conservation. Labeled residues were mutated in this study (Fig. 3). (E) Overlay of the apo and co-complex structures of *St*CphA2, showing the presence of substrates leads the $G_{lid}$ lobe to adopt a more closed conformation.

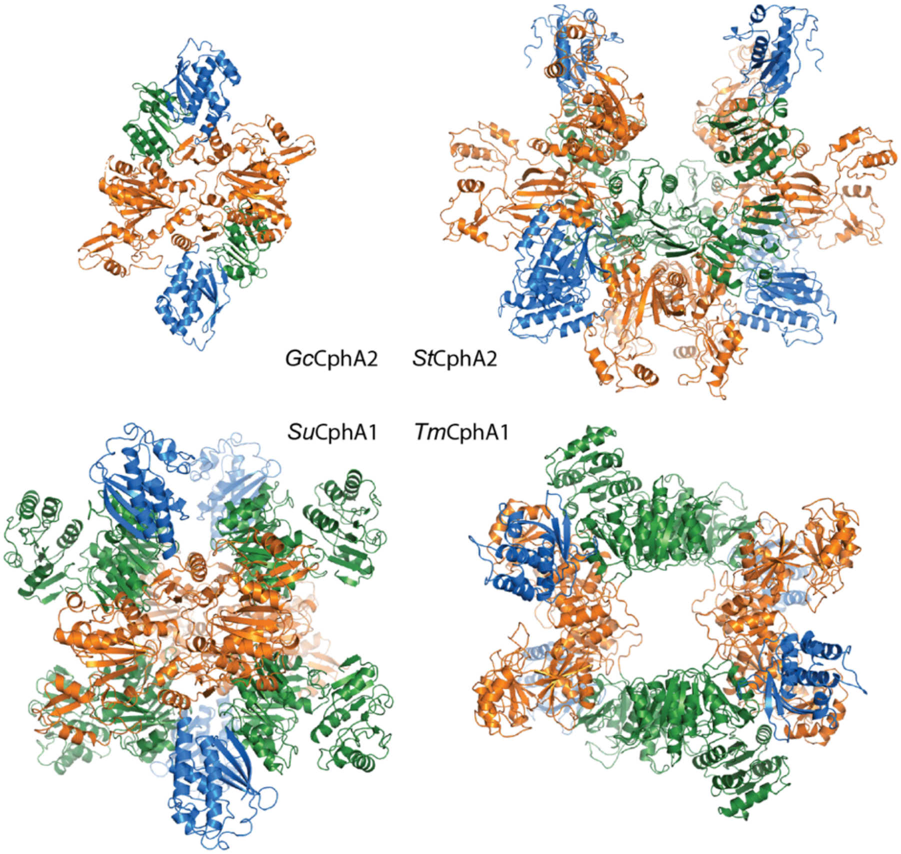

**Supplemental Figure 3.** Oligomeric architectures of cyanophycin synthetases. *G. citriformis* (*Gc*)CphA2) is dimeric(Sharon et al. 2022a), *Stanieria* sp. NIES-3757 (*St)*CphA2 is hexameric and *Synechocystis sp.* UTEX2470 (*Su*)CphA1 and *Tatumella morbirosei* DSM23827 (*Tm*)CphA1 are tetrameric(Sharon et al. 2021; Sharon et al. 2022b), as are *Acinetobacter baylyi* DSM587 (*Ab*)CphA1(Sharon et al. 2021) and *Trichodesmium erythraeum* (*Te*)CphA1(Miyakawa et al. 2022) (not shown).
